# Topological benchmarking of algorithms to infer Gene Regulatory Networks from Single-Cell RNA-seq Data

**DOI:** 10.1101/2022.10.31.514493

**Authors:** Niclas Popp, Marco Stock, Jonathan Fiorentino, Antonio Scialdone

## Abstract

In recent years, many algorithms for inferring gene regulatory networks from single-cell transcriptomic data have been published. Several studies have evaluated their accuracy in estimating the presence of an interaction between pairs of genes. However, these benchmarking analyses do not quantify the algorithms’ ability to capture structural properties of networks, which are fundamental, for example, for studying the robustness of a gene network to external perturbations. Here, we devise a three-step benchmarking pipeline called STREAMLINE that quantifies the ability of algorithms to capture topological properties of networks and identify hubs. To this aim, we use data simulated from different types of networks as well as experimental data from three different organisms. We apply our benchmarking pipeline to four algorithms and provide guidance on which algorithm should be used depending on the global network property of interest.

## 1 Introduction

Single-cell transcriptomics techniques allow probing patterns of gene expression on an increasingly larger scale, with recent studies including millions of cells and thousands of genes [1]. Such rapid progress in expanding the scale of available data makes single-cell datasets more appealing for tasks like the inference of gene regulatory networks (GRNs), with the goal of achieving a mechanistic understanding of the systems at hand and going beyond purely descriptive characterizations [2]–[4]. However, GRN inference from single-cell data entails many computational challenges, such as high levels of technical noise in the data [5], the extreme sparsity of the ground truth network to be inferred [6] and the increasing scale of gene expression data [7]. For this reason, many algorithms for GRN inference from single-cell data have been published in the last few years. The increasingly large number of such algorithms demands benchmarking studies that can guide the user in the choice of the best-performing methods under various conditions [8]–[10].

While the benchmarking studies that have been published offer some guidance for users, they are affected by important limitations. First, the quantification of the performance is obtained from a limited number and types of networks. Moreover, the available benchmarking studies mostly focus on the ability of the GRN algorithms to predict local features of networks, like the interactions between pairs of genes, using, for example, area under the curve metrics, or the presence of specific sub-graphs (network motifs). Nevertheless, these metrics do not assess the algorithms’ ability to infer the structural properties of the GRN, which quantify important features like the robustness to perturbations [11] and the presence of network hubs representing master regulators. The robustness of GRNs is one of their main characteristics [12], and for this reason, their topology is also studied to improve the robustness of general network structures in other fields, such as wireless sensor networks [13], [14]. Moreover, the inclusion of network topology in GRN inference methods has also been shown to improve their performance, e.g., using microarray and bulk RNA-seq data [15], [16]. Recently, an algorithm based on a global network centrality measure and local network motifs has been introduced [17], showing an improvement in inference performance due to the reduction of network redundancy. This class of inference methods is still lacking for single-cell RNA-seq data.

The structural properties of networks can be quantified by topological measurements [18], including, for instance, the network efficiency and the assortativity. So far, the performance of GRN inference algorithms on the estimation of topological properties has been only assessed with bulk RNA-seq data [19], [20], and employing a limited number of synthetic networks [20], which makes it hard to reach robust conclusions for single-cell data.

In this work, we developed STREAMLINE, a three-step benchmarking framework to score the performance of GRN inference algorithms in estimating structural properties of networks from single-cell RNA-seq (scRNA-seq) datasets. The structural properties we considered quantify the network’s robustness to perturbations and the presence of hubs. We used data simulated from hundreds of networks belonging to four classes with different structural properties [21], [22], as well as from a set of curated networks extracted from real GRNs [8]. In addition to simulated data, we also used real datasets from yeast, mouse, and human [23].

We applied STREAMLINE to four GRN inference algorithms chosen amongst the top-performing ones [8]. Our benchmarking analysis provides guidance in the choice of the algorithm for the prediction of network robustness and the identification of hubs. Moreover, our results point to systematic biases in some algorithms, which could indicate ways of improving them.

To facilitate the use of our benchmarking framework, we made it compatible with an existing pipeline (BEELINE [8]), and we made all the code available in a GitHub repository (github.com/ScialdoneLab/STREAMLINE).

## 2 Results

### 2.1 Overview of STREAMLINE

The steps involved in STREAMLINE are schematically represented in Figure 1. We consider two types of datasets: simulated and real datasets. With the simulated datasets, we generate scRNA-seq data in silico from four classes of networks with well-defined and different structural properties to be able to test the algorithms in different scenarios. The classes of networks we consider are Random, Small-World, Scale-Free, and Semi-Scale-Free Networks. Random or Erdös-Renyi (ER) networks include a set of nodes in which each node pair has the same probability of being connected by an edge [24]. We include this class of networks as a control. In Scale-Free (SF) networks, the edges are drawn such that the degree distribution follows a power law [25]. SF networks have been considered ubiquitous in cell biology [26], but their presence, at least on a global network level, is still debated [27]. For this reason, we also employ Semi-Scale-Free (SSF) networks, in which only the out-degree distribution follows a power law, while the in-degree distribution is uniform. Such networks were introduced by Ouma et al. [21] to model real GRNs. Small-World (SW) networks have the property that the neighbors of any given node are likely to be neighbors of each other [22]. The small-world property has been observed, for instance, in yeast [28] and human lung cancer [29] GRNs. In addition to these, we included four Curated (Cur) networks that consist of sub-networks of known GRNs [8]. Networks from each class are defined by a set of parameters. To make our results independent of specific instances of networks, we sampled 20 networks from each class with different combinations of parameters and two sizes: a smaller (15 nodes and 50 edges) and a larger (25 nodes and 100 edges) size. All results shown below are averaged over all the instances of networks generated for a given class. Details about the network classes and the parameters used for network sampling are provided in the Methods section, Supplementary Section S1 and Table S1. From each of these networks, we simulated scRNA-seq datasets using BoolODE, a recently developed software based on ordinary differential equations [8] (see Methods section).

**Figure 1:**
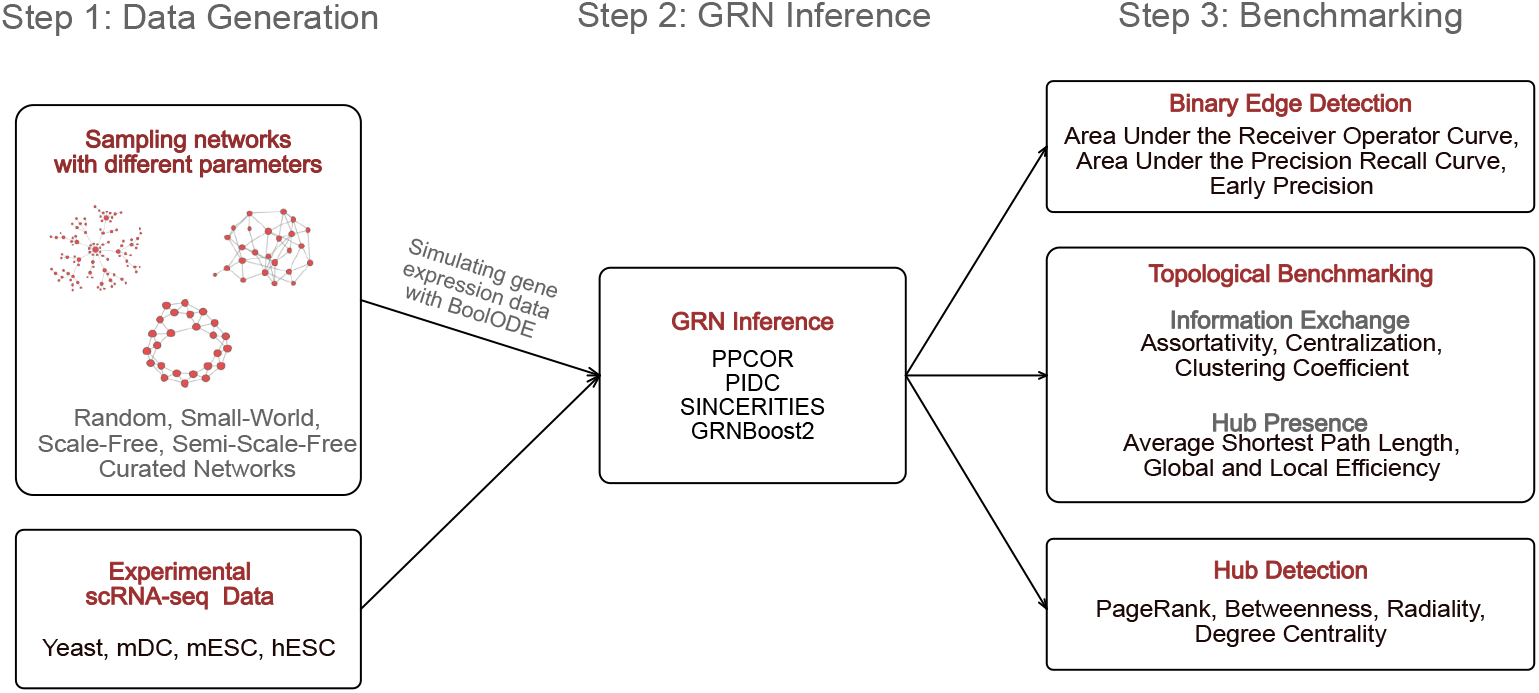
Schematic overview of STREAMLINE. STREAMLINE consists of three steps: first, synthetic scRNA-seq data are generated from different classes of networks. Then, GRN inference methods are applied to synthetic as well as real data. Finally, the methods’ performance on the predictions of edges and of structural network properties (quantifying the network robustness and hub presence) are evaluated.

In addition to simulated datasets, we also considered four real scRNA-seq datasets generated from different organisms and cell types: yeast [30], mouse dendritic cells (mDC) [31], mouse embryonic stem cells (mESC) [32] and human embryonic stem cells (hESC) [33]. These datasets were used in a previous benchmarking study [23], where the authors also provide estimations of ground truth networks. The second step of our pipeline involves running the algorithms to infer GRNs from each of the datasets. We chose the four top-performing algorithms according to a recent study where the accuracy in predicting gene-gene interactions was evaluated [8]: PIDC [34], PPCOR [35], SINCERITIES [36] and GRNBoost2 [37]. Two of these methods (PIDC and PPCOR) give as output undirected networks, while SINCERITIES and GRNBoost2 provide directed networks. A brief description of each algorithm is included in the Methods section.

In our analysis, we first scored each method’s ability to predict the presence of edges. Specifically, we calculated both the area under the receiver operator curve (AUROC) and the area under the precision-recall curve (AUPRC) on the synthetic data, as well as the early precision (EPr) on the experimental data. The results are then grouped for each network class or organism, and we found results in line with previous studies (see [8], [9], Supplementary Section S2 and Figure S1).

Then, we analyzed the ability of each method to predict global properties of networks. In particular, we computed topological properties that quantify how efficiently information is exchanged in the network and the tendency of networks to include hubs.

The efficiency of information exchange measures how the behavior of a network can change following variations in its topology due to, e.g., the failure of some of its constituents [38]. In this context, it is crucial to quantify and assess the stability of a GRN, as it is subject to random errors due to mutations and extreme conditions that can impede regulatory interactions [39]. These aspects can be captured by the following topological measures: the Global Efficiency, the Local Efficiency and the Average Shortest Path Length (see Methods). Efficiency measures have already been used to study the relationship between evolutionary and topological properties of human gene regulatory networks [40], while the Average Shortest Path Length has been widely adopted as a measure of biological network navigability (defined as the ability of efficiently moving from a source to a target node through short communication paths), which is crucial for information distribution [41], [42].

Another biologically important property of networks is the presence of hubs, i.e., nodes that have a degree much larger than the average. In GRNs, hubs are genes that regulate the expression levels of many other genes and can represent master regulators of a given biological process. Through structural network analysis, it has been shown that the presence of hubs is highly sensitive to perturbations in network topology [43], and it is linked to topological quantities like the Centralization, the Assortativity and the Clustering Coefficient [44], [45], which we compute in STREAMLINE.

In addition to quantifying the tendency of networks to possess hubs, it is important to identify them correctly. Hence, we tested the GRN inference algorithms for their ability to predict which nodes constitute hubs. To do so, we computed four metrics used to detect hubs [46] - Page Rank Centrality, Betweenness Centrality, Out-Centrality and Radiality (see Methods) - and we compared the values obtained from the ground truth networks versus those calculated from the inferred networks.

Below, we describe the detailed results of each of these benchmarking analyses.

### 2.2 Estimation of information exchange efficiency

In order to quantify topological aspects of the efficiency in information exchange, we evaluated the Average Shortest Path Length, the Global Efficiency, and the Local Efficiency (Figure 2A) of the inferred and ground truth networks. In particular, we estimated the Mean Signed Error (MSE) between these quantities computed on the ground truth networks and the networks inferred from each of the algorithms (see Methods section). This enables to quantify the accuracy when estimating efficiency and robustness for each network class separately.

**Figure 2:**
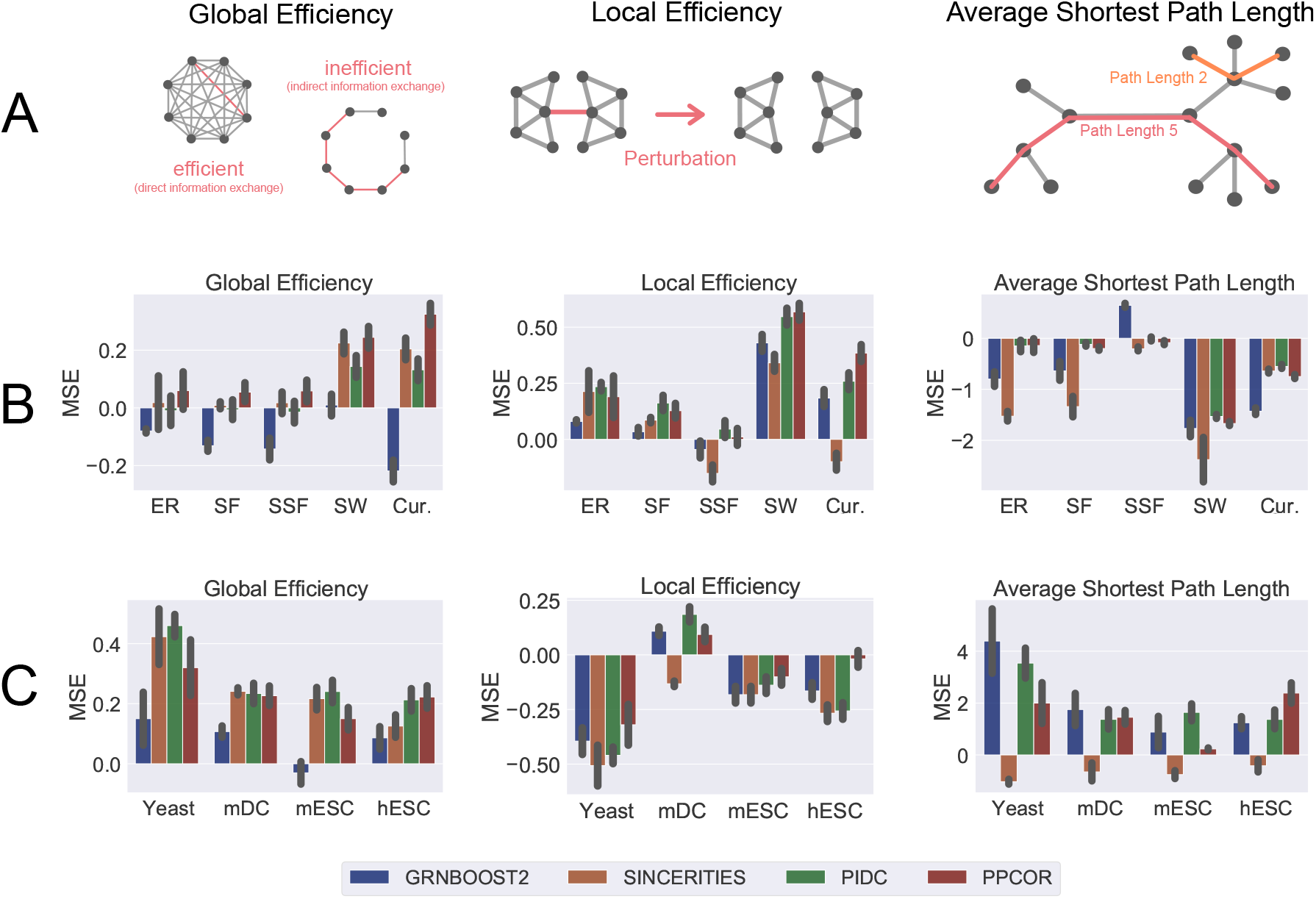
Results of the topological benchmarking of GRN inference algorithms with respect to information exchange both on synthetic and experimental scRNA-seq datasets. (A) Schematic representations of the three topological measures we computed (see Methods). Global Efficiency quantifies how well information can be distributed in the entire network. Local Efficiency measures how robust the network is to perturbation on a small scale. The Average Shortest Path Length specifies how many links are necessary to go from one node to another on average. (B) Barplots showing the Mean Signed Error (MSE) for the estimations of the topological properties written at the top in different types of synthetic networks (indicated on the x-axis) and for different algorithms (marked by colors). (C) Same as B, for networks estimated from real scRNA-seq datasets (indicated on the x-axis). The error bars display the standard deviations.

First, we considered the simulated datasets generated from different classes of networks. The different structural properties of each class of networks are reflected by different values of these topological measures, as shown in Figure S2A. For example, the SW networks are characterized by larger Average Shortest Path Length and lower Global and Local Efficiency, as expected based on their properties [22].

In Figure 2B, we report the MSE for all the topological measures computed on the simulated datasets. Overall, we found that the accuracy of the predictions depends on the type of network, in addition to the algorithm. For instance, all the tested algorithms tend to overestimate the Local Efficiency (MSE *>* 0), except for SINCERITIES, which underestimates it in SSF and Cur networks. The best predictions (corresponding to MSE∼ 0) are obtained on SF and SSF networks.

Similarly, the Global Efficiency tends to be overestimated by all algorithms except for GRNBoost2 (Figure 2B), which underestimates it. These results suggest the presence of biases that make the best-performing algorithm contingent on the type of network: for example, GRNBoost2 provides an accurate estimation of the Global Efficiency in SW networks, which is lower than in other network classes (Figure 2B and Figure S2).

The best estimations of the Average Shortest Path Length (Figure 2B) are provided by the PIDC and PPCOR algorithms, especially in the ER, SF, and SSF networks. In the SW and the Cur networks, for which the Average Shortest Path Length is greater than for SSF graphs (Figure S2A), all algorithms underestimate this property, particularly GRNBoost2 and SINCERITIES. This result contrasts the performance measured using statistical metrics where all algorithms apart from SINCERITIES perform best on SW networks (see Supplementary Section S2 and Figure S1).

We performed the same analysis on four real scRNA-seq datasets from three species (Figure 2C). The corresponding ground truth networks have lower Global and Local Efficiency and larger Average Shortest Path Length than the synthetic networks we considered (Figure S2B).

These differences likely affect the values of the MSE we computed, which are reported in Figure 2C. For example, unlike with the synthetic networks, we found an overall tendency of all algorithms to underestimate the Local Efficiency and overestimate the Global Efficiency. GRNBoost2 and PPCOR provide the most accurate predictions of the Global and Local Efficiency, respectively.

As for the Average Shortest Path Length, the MSE is mostly positive, indicating an overestimation, and it is smallest for SINCERITIES, which is the best-performing algorithm in this case.

### 2.3 Hub analysis

One important downstream analysis on GRNs is the identification of genes with a number of links much larger than the average. These are known as network hubs, which can play key roles in differentiation and reprogramming [47] and have been identified as potential disease regulators or drug targets [48]. The presence of hubs depends on several topological properties that are influenced by the type of network. In particular, we expect the hubs in SF and SSF networks to be more easily identifiable due to their node degree distribution. This is reflected, for example, by the higher Centralization values of SF and SSF networks compared to other classes of networks (Figure S2).

Here, we first analyzed how well each algorithm predicts the value of topological measures that quantify the tendency of networks to include hubs. Then, we test how well the inferred GRN preserves hub identities.

#### 2.3.1 Hub-related topological quantities

The topological measures we chose to quantify the presence of hubs are Assortativity, Clustering Coefficient and Centralization (Figure 3A). In networks with larger values of Assortativity, nodes with lower degrees tend to be linked to nodes that feature a higher degree; hence, in these networks, hubs tend to be present and clearly identifiable. Networks with a large Clustering Coefficient feature groups of nodes with high interconnectivity that, thus, have similar node degrees. In this situation, hubs are less dispersed. The Centralization quantifies how centralized a graph is around a small number of nodes, which will have a large number of links, and will therefore tend to be strong and clearly identifiable hubs.

**Figure 3:**
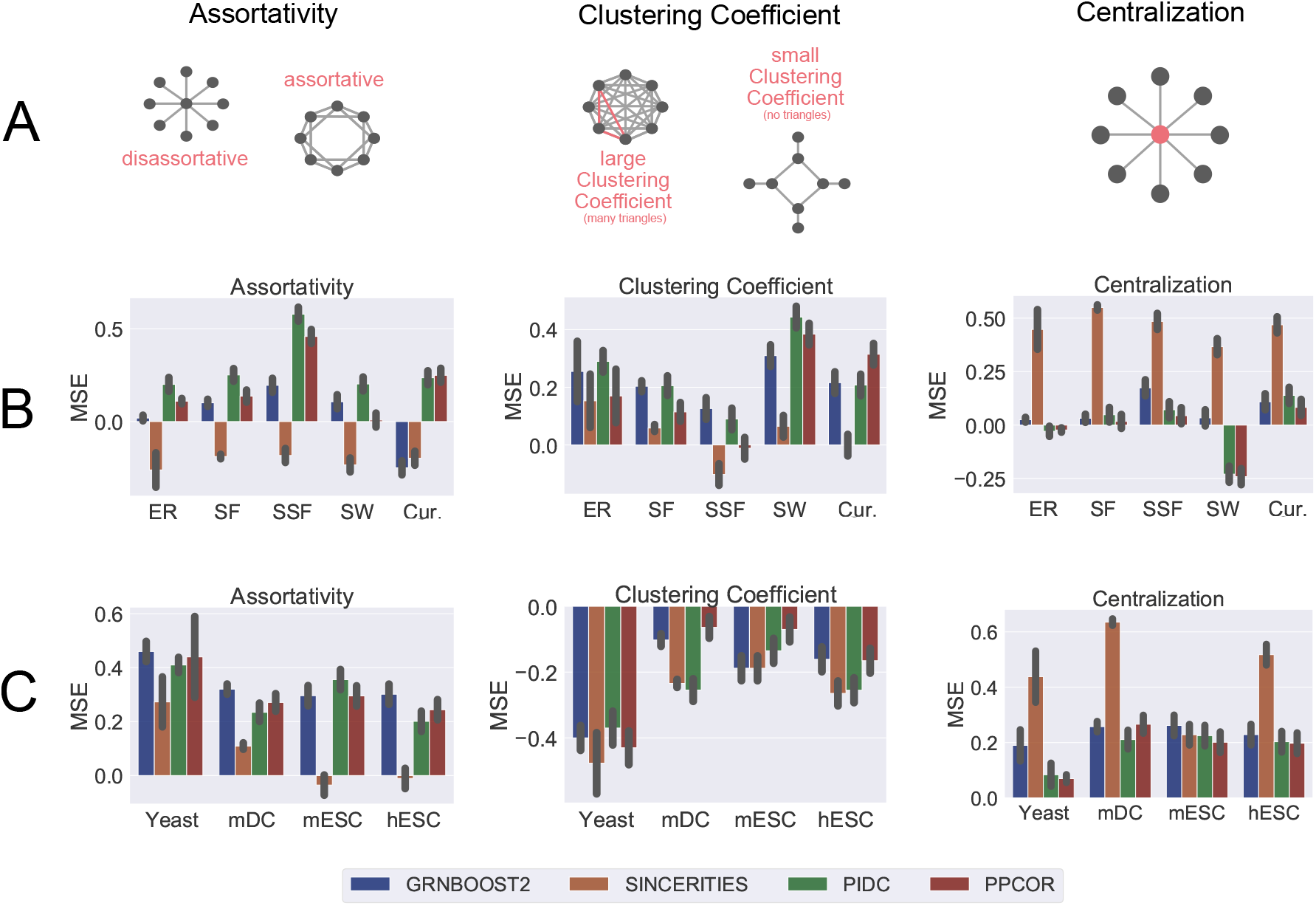
Results for the topological benchmarking of GRN inference assessing the presence of hubs. (A) Schematic representation of the three topological measures considered here (see Methods). The Assortativity quantifies the tendency of nodes in the networks to attach to others with similar degrees. The Clustering Coefficient reflects how much the nodes in a graph tend to cluster together. The Centralization indicates how strongly the network is arranged around a single center. (B) Barplots showing the Mean Signed Error (MSE) for the estimations of the topological properties written at the top in different types of synthetic networks (indicated on the x-axis) and for different algorithms (marked by colors). (C) Same as B, for networks estimated from real scRNA-seq datasets (indicated on the x-axis). The error bars display the standard deviations.

The networks we simulated data from are characterized by different values of the above topological measures due to their properties (see [21] and Figure S2A). This allowed us to explore the performance of the algorithms under different conditions. The three network properties assessed during this step uncovered multiple algorithm-specific behaviours (Figure 3B). The most evident involves the algorithm SINCERITIES, which yielded GRNs with lower Assortativity and larger Centralization than the corresponding ground truth graphs. On the other hand, the estimations of the Clustering Coefficient obtained with SINCERITIES tend to be more accurate than those coming from other algorithms.

As in the previous section, we repeated the analysis on real datasets (Figure 3C). The corresponding networks have Assortativity values that are lower than those of most synthetic networks and are in line with the scale-free hypothesis for GRNs [21]. On average, the Clustering Coefficients are similar to those of ER or SW networks (Figure S2B). Such values of the topological properties indicate that these networks display a higher tendency to contain hubs that are more clustered together when compared to random networks.

Similarly to what happens with the synthetic datasets, here, the Centralization is overestimated by SINCERITIES (Figure 3C). In contrast, the Assortativity is now preserved better by SINCERITIES rather than being underestimated.

GRNBoost2, PIDC and PPCOR show similar performances. These three algorithms overestimate the Assortativity as well as the Centralization, albeit less than SINCERITIES. Furthermore, the Clustering Coefficient is underestimated by all algorithms.

The difference in performance between the real and the synthetic data might be due to a number of factors. First, the gold standards provide only estimates for GRNs of the three organisms, while the synthetic data are simulated from fully specified networks. Furthermore, we found that the algorithms tend to output networks that feature similar topological properties, regardless of the data they are run on. This might explain, for example, the opposite trends in the estimations of the Assortativity and the Clustering Coefficient with the synthetic versus the real datasets.

#### 2.3.2 Hub identification

While hubs are loosely defined as nodes having degrees higher than average, there is no consensus on the best metric to identify them. For this reason, here we compute four centrality measures that have been previously adopted to find hubs in GRNs [46]: the Betweenness [49], the Out-Centrality [50], the Radiality [51], and the Page Rank [52] (see Methods). Among these, the Page Rank and the Out-Centrality metrics are conserved along the evolution and relevant in pluripotent cells [53]. Moreover, they were proposed as metrics to distinguish life-essential versus specialized subsystems [53].

We verified how accurately the hub-identification measures are estimated by the four inference algorithms introduced above. More specifically, we selected the set of top 10% nodes according to the centrality measure computed in the ground truth network, Ω_*true*_, and in the inferred network, Ω_*inferred*_. Then, we quantified the similarity between the two sets of nodes with the Jaccard similarity index [54] (see Methods). Finally, we computed the ratio between *J* and *J*_*rand*_, i.e., the Jaccard index between Ω_*true*_ and a set of randomly selected nodes Ω_*rand*_ (see Methods). Hence, the ratio *J/J*_*rand*_ shown in Figure 4 represents the improvement in the prediction of hubs in the inferred network compared to a random guess.

**Figure 4:**
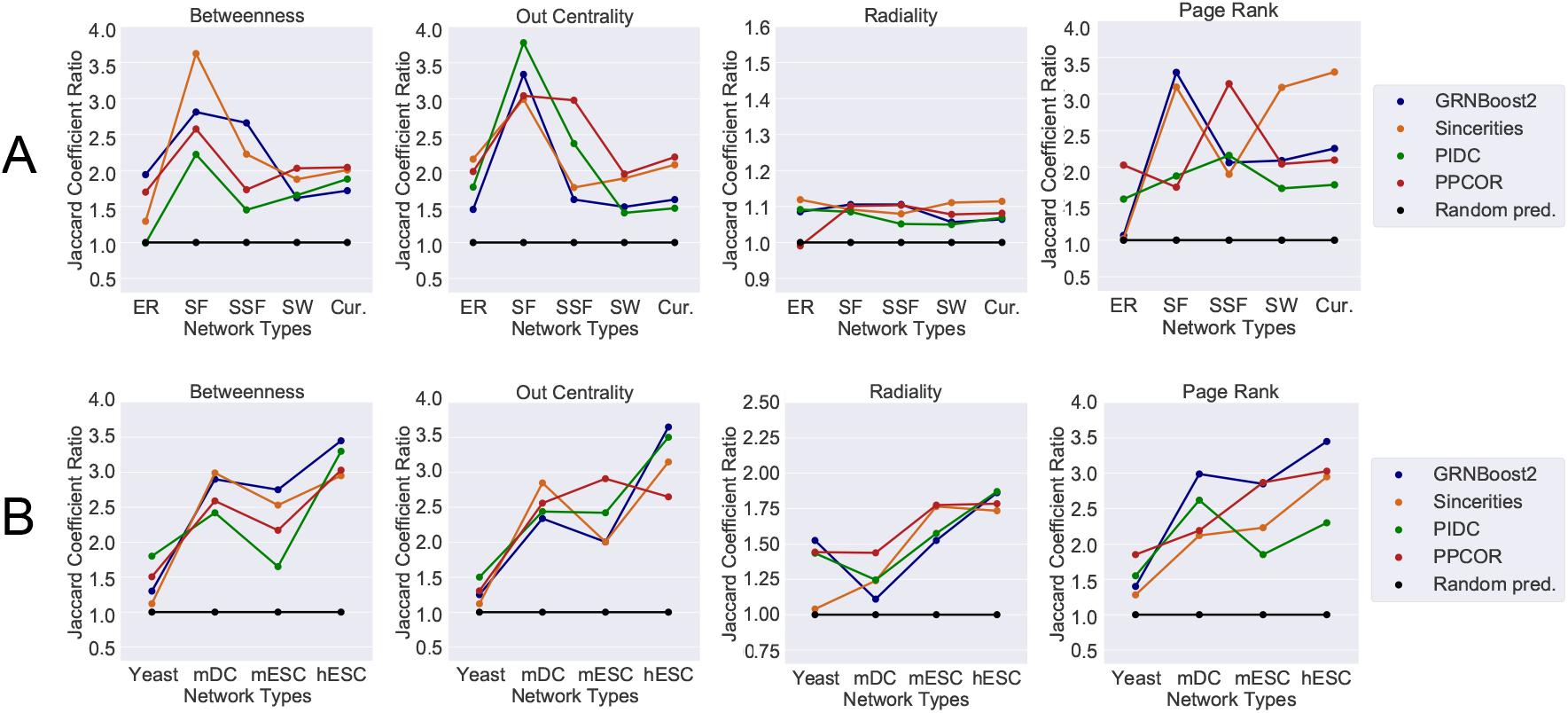
Accuracy of hub detection. The accuracy is measured by the Jaccard coefficient ratio with a random predictor as reference (see Methods and Eq. 15). (A) The Jaccard coefficient ratio is plotted with various hub metrics (written on top) as a function of the type of synthetic network indicated on the x-axes for different algorithms. (B) Same as (A), plotted as a function of the networks inferred from real scRNA-seq datasets (indicated on the x-axes). The Betweenness estimates the influence that a node has on the information exchange in a graph based on path lengths. The Out Centrality is the out-degree of a node in directed networks or its overall degree in undirected networks. The Radiality assigns high centrality values to nodes with a short distance to all vertices in their reachable neighborhood compared to the graph diameter. PageRank is a generalization of the degree centrality that considers the eigenvalues of a modified adjacency matrix. We provide a detailed definition of the hub metrics in the text and the Methods section.

Generally, we find that better scores are achieved on networks featuring stronger hubs (i.e., with larger values of Centralization, like the GRN associated with hESC; see Figure S2). However, the scores depend on the algorithm as well as the network type. In real networks, there is a wide range of Centralization values, with the hESC networks featuring higher values, which indicate the presence of more clearly identifiable hubs. Conversely, the yeast GRN has lower values of Centralization and, thus, fewer hubs or hubs that are less isolated (Figure S2). As a possible consequence, the best performance is achieved on hESC networks. Similarly, the performance on synthetic data is best on SF networks which are likely to possess hubs that are easier to identify due to their degree distribution.

## 3 Discussion

Here, we performed benchmarking analyses to evaluate how well GRN inference algorithms can estimate the structural properties of the networks. More specifically, we quantify the ability of the algorithms to infer their robustness to perturbations as well as the presence and identification of network hubs. For this purpose, we computed six topological measures and tested four metrics for hub identification that are widely known and used in network theory. Moreover, we considered scRNA-seq data simulated from different types of networks as well as real data collected from different organisms.

In this extensive benchmarking, we focused on network properties (i.e., robustness and hubs) that are not taken into account in currently available benchmarking studies performed on scRNA-seq data, despite being considered important when studying GRNs and, more in general, biological networks (see, e.g., [12], [55], [56]). For example, the identification of putative master regulators via degree-based measures on GRNs is a very commonly used practice (see, e.g., [46], [57], [58]).

The benchmarking results are summarized in Table 1.

**Table 1:**
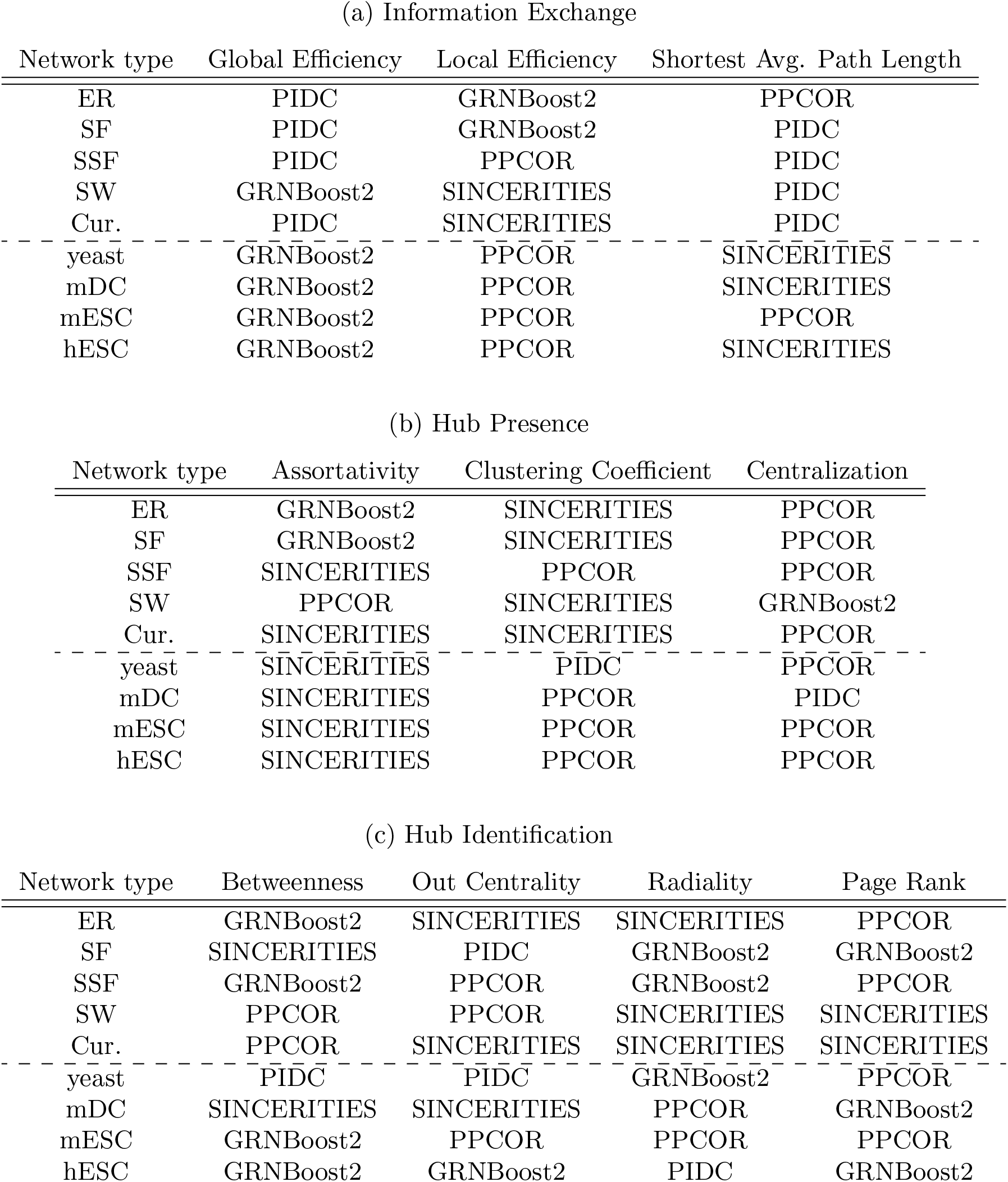
The above table summarizes which algorithms performed best in each step of our benchmarking pipeline, subdivided by dataset and property type. The top-performing methods were selected according to the lowest Mean Signed Error (MSE) in absolute value.

| Network type | Global Efficiency | Local Efficiency | Shortest Avg. Path Length |
| --- | --- | --- | --- |
| ER | PIDC | GRNBoost2 | PPCOR |
| SF | PIDC | GRNBoost2 | PIDC |
| SSF | PIDC | PPCOR | PIDC |
| SW | GRNBoost2 | SINCERITIES | PIDC |
| Cur. | PIDC | SINCERITIES | PIDC |
| yeast | GRNBoost2 | PPCOR | SINCERITIES |
| mDC | GRNBoost2 | PPCOR | SINCERITIES |
| mESC | GRNBoost2 | PPCOR | PPCOR |
| hESC | GRNBoost2 | PPCOR | SINCERITIES |

(b) Hub Presence
| Network type | Assortativity | Clustering Coefficient | Centralization |
| --- | --- | --- | --- |
| ER | GRNBoost2 | SINCERITIES | PPCOR |
| SF | GRNBoost2 | SINCERITIES | PPCOR |
| SSF | SINCERITIES | PPCOR | PPCOR |
| SW | PPCOR | SINCERITIES | GRNBoost2 |
| Cur. | SINCERITIES | SINCERITIES | PPCOR |
| yeast | SINCERITIES | PIDC | PPCOR |
| mDC | SINCERITIES | PPCOR | PIDC |
| mESC | SINCERITIES | PPCOR | PPCOR |
| hESC | SINCERITIES | PPCOR | PPCOR |

| Network type | Betweenness | Out Centrality | Radiality | Page Rank |
| --- | --- | --- | --- | --- |
| ER | GRNBoost2 | SINCERITIES | SINCERITIES | PPCOR |
| SF | SINCERITIES | PIDC | GRNBoost2 | GRNBoost2 |
| SSF | GRNBoost2 | PPCOR | GRNBoost2 | PPCOR |
| SW | PPCOR | PPCOR | SINCERITIES | SINCERITIES |
| Cur. | PPCOR | SINCERITIES | SINCERITIES | SINCERITIES |
| yeast | PIDC | PIDC | GRNBoost2 | PPCOR |
| mDC | SINCERITIES | SINCERITIES | PPCOR | GRNBoost2 |
| mESC | GRNBoost2 | PPCOR | PPCOR | PPCOR |
| hESC | GRNBoost2 | GRNBoost2 | PIDC | GRNBoost2 |

First, we observe that the algorithms’ performance in edge detection (Figure S1) does not correlate with their performance in estimating the topological properties, which proves the need for a targeted benchmarking analysis like ours. Overall, we found that there is no single best-performing algorithm, but the performance depends on the properties of the ground truth network and the topological metric being considered. For example, with real datasets, GRNBoost2 achieves the best results in estimating Global Efficiency; whereas SINCERITIES produces the most accurate estimations of Local Efficiency and Average Shortest Path Length in almost all the real datasets (Figure 2). A similar situation emerged for the metrics quantifying Hub Presence (Figure 3), where the estimations of Assortativity obtained with SINCERITIES are the most accurate, while PPCOR performed best when estimating the Clustering Coefficient and the Centralization in almost all the real datasets.

The situation for the Hub Identification is more complex, as the best-performing algorithm seems more susceptible to the specific dataset analyzed (Figure 4). However, overall, all algorithms achieve good performance with all datasets and metrics. A notable exception to this is the yeast dataset (Figure 4B), where the accuracy of predictions for all algorithms (and especially SINCERITIES) was relatively low. This might be linked to the ground truth network of yeast having less clearly defined hubs, as, e.g., the value of Centralization suggests (Figure S2).

The benchmarking done with synthetic networks allowed us to check the performance of algorithms with networks having specific and tunable properties. In some cases, this has brought to light specific biases present in the networks estimated by each algorithm. For example, SINCERITIES produces more disassortative and centralized networks (i.e., networks with relatively low Assortativity and high Centralization), which causes an underestimation of Assortativity and overestimation of Centralization for all types of synthetic networks (Figure 3). Similar observations can be made, for example, with GRNBoost2, which tends to generate networks with lower Global Efficiency (Figure 2).

The algorithm biases that we detected are likely to derive from the specific strategies that each algorithm follows to estimate GRNs. Importantly, the knowledge of such biases can guide the effort to improve current algorithms, for example, by assisting in the design of objective functions that can lead to networks with global properties closer to real GRNs. This approach can be justified by the observation that GRNs share certain topological features, such as a scale-free [59] or semi-scale free [21] node-degree distributions, which could be assumed as prior knowledge during the inference process.

Finally, the topological quantities we considered can also be used to optimize community-based inference schemes. Currently, consensus networks are derived from the outputs of different methods by taking into account only their performance in estimating single edges. Instead, new strategies could be devised that also consider the estimation of the network’s global properties.

## 4 Methods

### 4.1 Ground Truth networks and Gold Standards

#### 4.1.1 Synthetic networks

We use parameter-controlled networks from four different classes as well as the Curated GRNs that have been used in BEELINE [8]. The output of the network samplers is a graph *G* with *n* nodes and *m* edges.

##### Random networks

Random networks were created by the Erdös-Renyi *G*(*n, p*) model, which outputs a graph with *n* nodes where each pair is connected with probability *p* [24]. We set *p* so that the expected number of edges equals *m*.

##### Scale-Free networks

Networks with a degree distribution that follows a power law are classified as Scale-Free [60]. Given the parameter *α*, the expected degree distribution follows

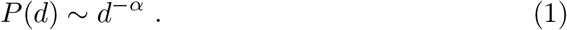

For directed networks the in-degree distribution and the out-degree distribution can feature different parameters *α*_*in*_ and *α*_*out*_. We applied combinations of different in- and out degrees. The exact values can be found in Table S1.

##### Semi-Scale-Free networks

Following the analysis of the degree distributions in known GRNs [21], we sample Semi-Scale-Free networks which feature an out-degree distribution that follows a power-law but a uniform in-degree distribution. Additionally, only 50% of the nodes have outgoing edges.

##### Small-World networks

We use the Watts-Strogatz model to sample networks that feature Small-World topology [22]. The algorithm starts with *n* nodes with degree *k* in a regular lattice and then rewires edges with probability *p*.

##### Curated networks

Curated networks are four known GRNs that were used in BEELINE to evaluate the statistical performance of the GRN inference algorithms [8]. These networks are simple models for mammalian cortical area development, ventral spinal cord development, hematopoietic stem cell differentiation and gonadal sex determination. We include them for comparison.

##### Network sampling

We use the Julia package LightGraphs.jl [61] to sample the networks explained above. The parameters were chosen such that a large variety of structurally different networks is covered. We simulate single-cell RNA-sequencing data using BoolODE [8]. For every parameter set we sampled data from 100 cells for 10 smaller and 10 larger networks. The exact specifications and number of networks can be found in Table S1.

#### 4.1.2 Experimental networks

For the benchmarking of GRN inference on experimental single-cell RNA-sequencing data we select four datasets from human [33], mouse [31], [32], and yeast [30] and compare the output networks to different types of gold standard networks that were collected by Stone et al. [23]. The properties of the networks and the number of corresponding gold standards can be found in Table S2. For the detailed description of the preprocessing we refer to [23].

### 4.2 Inference algorithms

We select the four top-performing algorithms from BEELINE [8] and examine the results using our three-step benchmarking pipeline.

#### GRNBoost2

GRNBoost2 [37] infers a GRN independently for each gene, by identifying the most important regulators using a regression model. It is an alternative to GENIE3, which uses a similar inference scheme but does not scale to larger datasets due to its runtime.

#### SINCERITIES

SINCERITIES [36] is a causality based method, that computes temporal changes in the expression of each gene. The GRN is inferred by solving a specifically formulated ridge regression problem.

#### PIDC

The PIDC inference scheme [34] is based on partial information decomposition, which is a multivariate information theoretic measure for triplets of random variables. Since it is symmetric, the resulting network is undirected.

#### PPCOR

PPCOR [35] calculates the partial and semi-partial correlation coefficients for every possible pair of genes. Edges are ranked according to these values. By using the correlation as sign, it is possible to assign a direction to the interactions but we found better topological performance in the undirected version.

### 4.3 Quantities used for benchmarking

#### 4.3.1 Binary Edge Detection

In order to statistically benchmark the binary classification problem of correctly retrieving edges we followed previous studies [8], [9] by calculating the area under the precision recall curve (AUPRC) and the area under the receiver operator curve (AUROC) on synthetic data. For the experimental datasets we use the early precision (EPr) among the top k edges, where k is the number of interactions in the corresponding gold standard. The EPr is better suited to classification accuracy on large datasets where the reference networks is not the entire ground truth.

#### 4.3.2 Topological graph properties related to information exchange

For our topological analysis we select six different properties that allow for a meaningful structural characterisation of a network and group them with respect to their matter as either being related to the exchange of information or the presence of hubs. We assume a graph with *n* nodes and *m* edges.

##### Average shortest path length

The *Average Shortest Path Length* measures by how many links two random nodes are connected on average. *d*(*v, w*) denotes the distance between two nodes *v* and *w*.

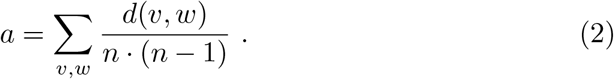

##### Global efficiency

The *Global efficiency* estimates how efficiently information is exchanged in the network on a global scale which relates this the concept of vulnerability of networks to the decrease in networks efficiency in the case when some of the components malfunction [38], [39]. It is given by

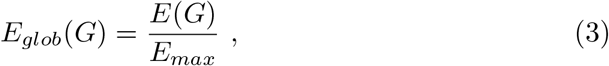

Where

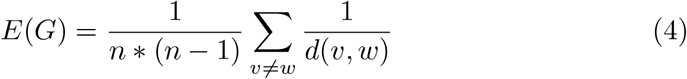

and *E*_*max*_ is the maximum *E*_*glob*_ for a fully connected graph given a number of nodes. Since gene regulation can be interpreted as information exchange between nodes in GRN, *E*_*glob*_ is a meaningful quantity to estimate. A more detailed explanation of the relation between global efficiency and vulnerability can be found in [62].

##### Local efficiency

The *Local efficiency* describes the resistance of the network to perturbation on a small scale [38]. Similarly to the global efficiency it is motivated by the behavior of the network when some of its constituents fail. A more detailed derivation can be found in [62]. The quantity is defined as

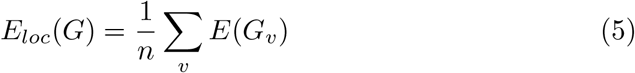

where *G*_*v*_ is the graph that only consists of *v* and its immediate neighbors. In practice there can be many factors that can perturb gene regulation, thus the local efficiency is an important score to be preserved, especially on experimental data.

#### 4.3.3 Topological graph properties related to the presence of hubs

##### Assortativity

The preference for a network’s nodes to attach to others that have a similar degree is captured by the Assortativity [63]. It is quantified by the *Assortativity Coefficient r*, which is the Pearson correlation coefficient between the degree *k* of a node and the average degree of its neighbors ⟨*k*_*nn*_⟩ when treating them as random variables.

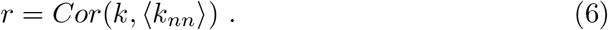

Networks with a negative Assortativity Coefficient are called disassortative, networks with a positive *r* are called assortative. Disassortative networks have a higher tendency to possess hubs, which is an important feature of GRN that we examine in section 2.3.

##### Centralization

The goal of the *Centralization H* is to provide an estimate of how centralized a graph is around the node *v*^*∗*^ which has the highest degree [50]. It is defined as

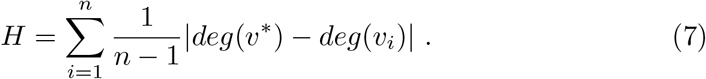

A highly centralized network is focused around a small number of nodes which could be identified as biologically important.

##### Clustering Coefficient

The Clustering Coefficient measures the extend to which nodes in a graph tend to cluster together. It is quantified by the *local clustering coefficient:*

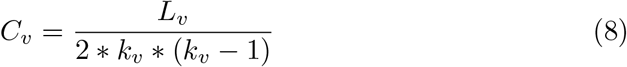

and the *global clustering coefficient* [22]:

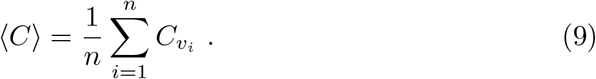

where *L*_*v*_ represents the number of links between the *k*_*v*_ neighbors of node *v*. For our analysis we focused on the global clustering coefficient since it captures clustering on a global scale. A network with a larger clustering coefficient is more interconnected which can result in more complicated gene regulations.

#### 4.3.4 Evaluation score

Since we are particularly interested in whether certain topological features are over- or underestimated we employ the Mean Signed Error (MSE) as evaluation quantity. For a property *P* which is being analyzed on networks *G*_1_, *G*_2_, …, *G*_*n*_ it is computed by

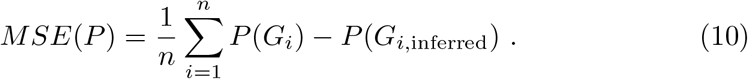

#### 4.3.5 Hub identification

##### Degree centrality

The degree centrality purely evaluates the degree of a node in a network. In a directed graph there are two different degree centralities: the in-degree centrality and the out-degree centrality. In the directed case we use the out-degree centrality, in the undirected case we use the overall degree centrality.

##### Betweenness centrality

Betweenness centrality describes the extend to which nodes stand between each other. It was formally defined by Freeman [49] as

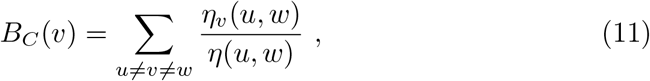

where *η*(*u, w*) is the number of shortest paths from *u* to *w* and *η*_*v*_(*u, w*) is the number of shortest paths from *u* to *w* passing through v. If the graph is not connected the betweenness centrality is evaluated for each connected component.

##### Page Rank centrality

The Page Rank centrality is the output of the Page Rank algorithm which is focused on link analysis. The output is a distribution that models the likelihood to reach any particular node when randomly moving along edges. Details of the algorithm can be found in [52].

##### Radiality

Radiality [51] considers the global structure of the networks and indicates how connected an individual is in the entire network structure. The definition makes use of the reverse distance matrix

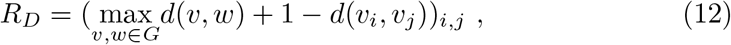

where *d*(*v, w*) describes the shortest path length from *v* to *w*. Radiality is defined as

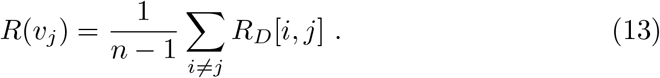

##### Evaluation score

We rank the nodes of each network according to the centrality measures defined above and in networks with more than 50 nodes we classify the top 10% as hubs while in smaller networks we select the top 20%. Our goal is to analyze the set similarity between the hubs in the ground truth or gold standard Ω_*true*_ and the inferred networks Ω_*inf*_. Therefore we compute the average Jaccard coefficient [54] for every network class, which is given by:

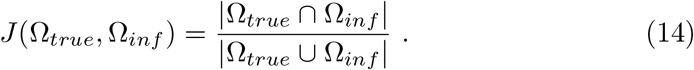

To quantify the performance of the different centrality measures, we use as evaluation score the ratio between the Jaccard coefficient based on the inferred networks and the expected coefficient for a random predictor which can be calculated explicitly [64]. If the fraction of nodes that is classified as hubs is given by *p* ∈ (0.1], the expected value for this Jaccard coefficient is given by:

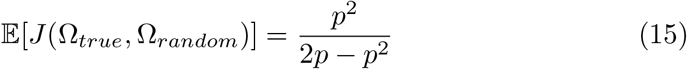

## Code availability

STREAMLINE is available at github.com/ScialdoneLab/STREAMLINE.

## Data availability

We used publicly available experimental single-cell RNA-sequencing data from yeast [30], mouse [31], [32] and human [33]. The gold standard networks were obtained from [23]. Single-cell RNA-sequencing data and the corresponding gold standards for Curated networks were obtained from [8]. The synthetic networks can be generated using the code provided in our Github repository github.com/ ScialdoneLab/STREAMLINE and the parameters listed in Supplementary Table S1, the corresponding single-cell RNA-sequencing data can be simulated using BoolODE [8].

## Acknowledgments

Jonathan Fiorentino and Marco Stock were supported by a Joachim Herz Stiftung Add-on Fellowship for Interdisciplinary Life Science. Marco Stock was supported by the Helmholtz Association under the joint research school “Munich School for Data Science - MUDS”. Work in the Scialdone lab is funded by the Helmholtz Association.

## Conflict of interests

The authors declare that they have no conflict of interest.

## Contributions

N.P. generated, analyzed data, produced the figures and interpreted the results.

M.S. provided advice and interpreted the results. J.F. and A.S. conceived the study and provided project supervision. N.P., M.S., J.F., A.S. wrote the manuscript.

## Supplementary Material

### S1. Parameters of the reference networks

The following two tables summarize the properties of the ground truth and gold standard networks. For the synthetic networks we list the parameters that were used for sampling as well as the number of networks that resulted. For the experimental networks the number of cells and genes are shown as well as the number of reference databases that were used. A more elaborated discussion of the properties of the real datasets is given by Stone et al.[23] who collected them.

**Table S1:**
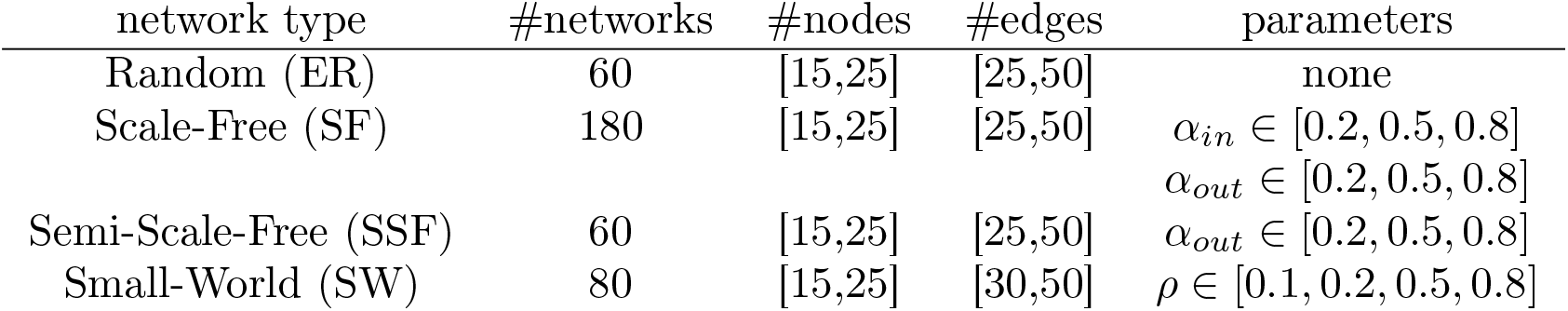
Parameters used to sample synthetic networks. We used all possible combinations which resulted in over 300 sampled networks in total.

**Table S2:** Properties of the experimental datasets and the number of reference databases used as gold standards.

| cell type | #cells | #genes | #reference databases |
| --- | --- | --- | --- |
| yeast [30] | 163 | 3847 | 7 |
| mDC [31] | 1211 | 9411 | 3 |
| mESC [32] | 2369 | 6618 | 10 |
| hESC [33] | 5520 | 7468 | 8 |

### S2. Binary Edge Detection

The results of our statistical benchmarking, shown in supplementary figure S1, reproduce the previous observation that inference accuracy in terms of AUPRC, AUROC and EPr is moderate at best [8], [9]. For synthetic data (Fig S1A-B), the highest prediction accuracy was achieved on Curated and Small-World networks, while on Scale-Free, Semi-Scale-Free and Random networks the performance is worse. Regarding the performance of the specific algorithms, GRNBoost2, PIDC and PPCOR perform similarly, while SINCERITIES identifies fewer correct interactions. This trend transfers to experimental data (Fig S1C). Additionally, the statistical performance on the yeast dataset is worse than on mDC, mESC and hESC.

**Figure S1:**
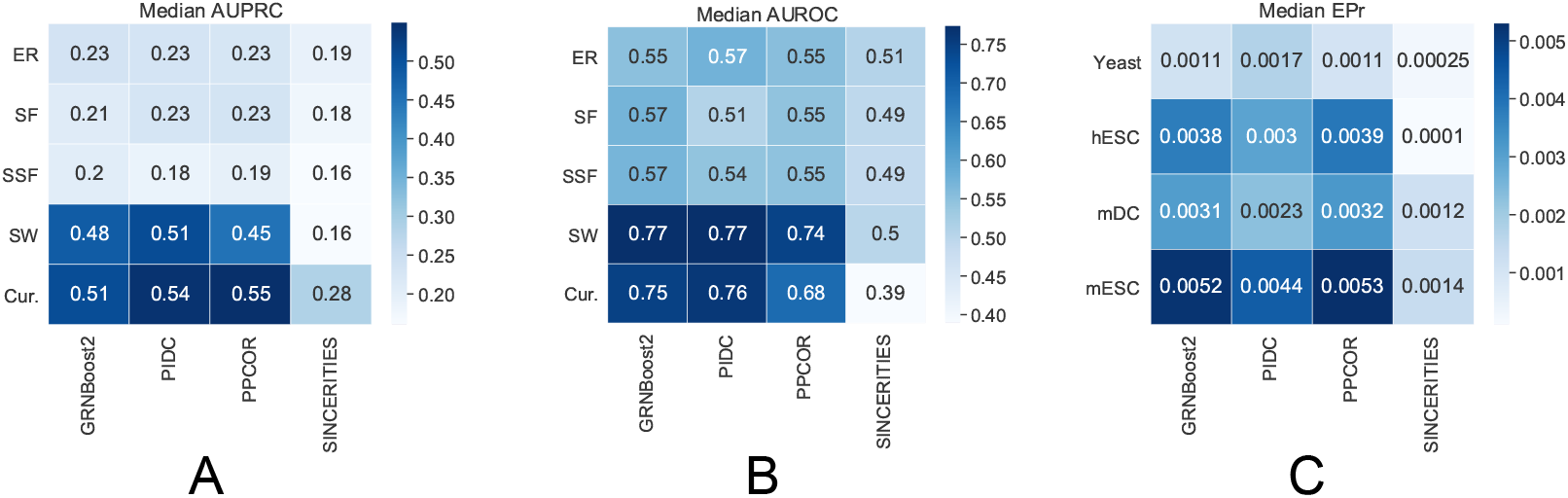
Benchmarking of edge detection of GRN inference algorithms on synthetic and experimental scRNA-seq datasets. The heatmaps report the median AUPRC and AUROC for synthetic data (panels A and B, respectively) and the median EPr for experimental data (panel C). Rows and columns correspond to GRN inference algorithms and network types, respectively.

**Table S3:** The table above lists the best performing algorithms for the task of edge detection from the ground truth networks as a binary classification problem. The best performing algorithms were selected according to the highest EPr scores for the experimental datasets or the highest AUPRC and AUROC scores for the synthetic datasets.

| Binary Edge Detection |  |  |  |
| --- | --- | --- | --- |
| Network type | EPr | AUPRC | AUROC |
| ER |  | PIDC | PIDC |
| SF |  | PPCOR | GRNBoost2 |
| SSF |  | GRNBoost2 | GRNBoost2 |
| SW |  | PIDC | GRNBoost2 |
| Cur. |  | PPCOR | PIDC |
| yeast | PIDC |  |  |
| mDC | PPCOR |  |  |
| mESC | PPCOR |  |  |
| hESC | PPCOR |  |  |

### S3. Topological properties of the reference networks

Figure S2 depicts the topological properties of the sampled ground truth networks as well as the gold standards for the experimental datasets. Among the networks we simulated data from, the Assortativity of SW and ER networks is relatively closer to 0 on average compared to other networks, while their Clustering Coefficient is lowest. Conversely, SF and SSF have lower (negative) Assortativity and larger Clustering Coefficients together with the Cur networks. This is in agreement with the known properties of each class of networks [21]. As previously mentioned, the Cur networks are a group of four small subnetworks of real GRNs, therefore their properties are influenced by their small size and individual peculiarities rather than general characteristics.

**Figure S2:**
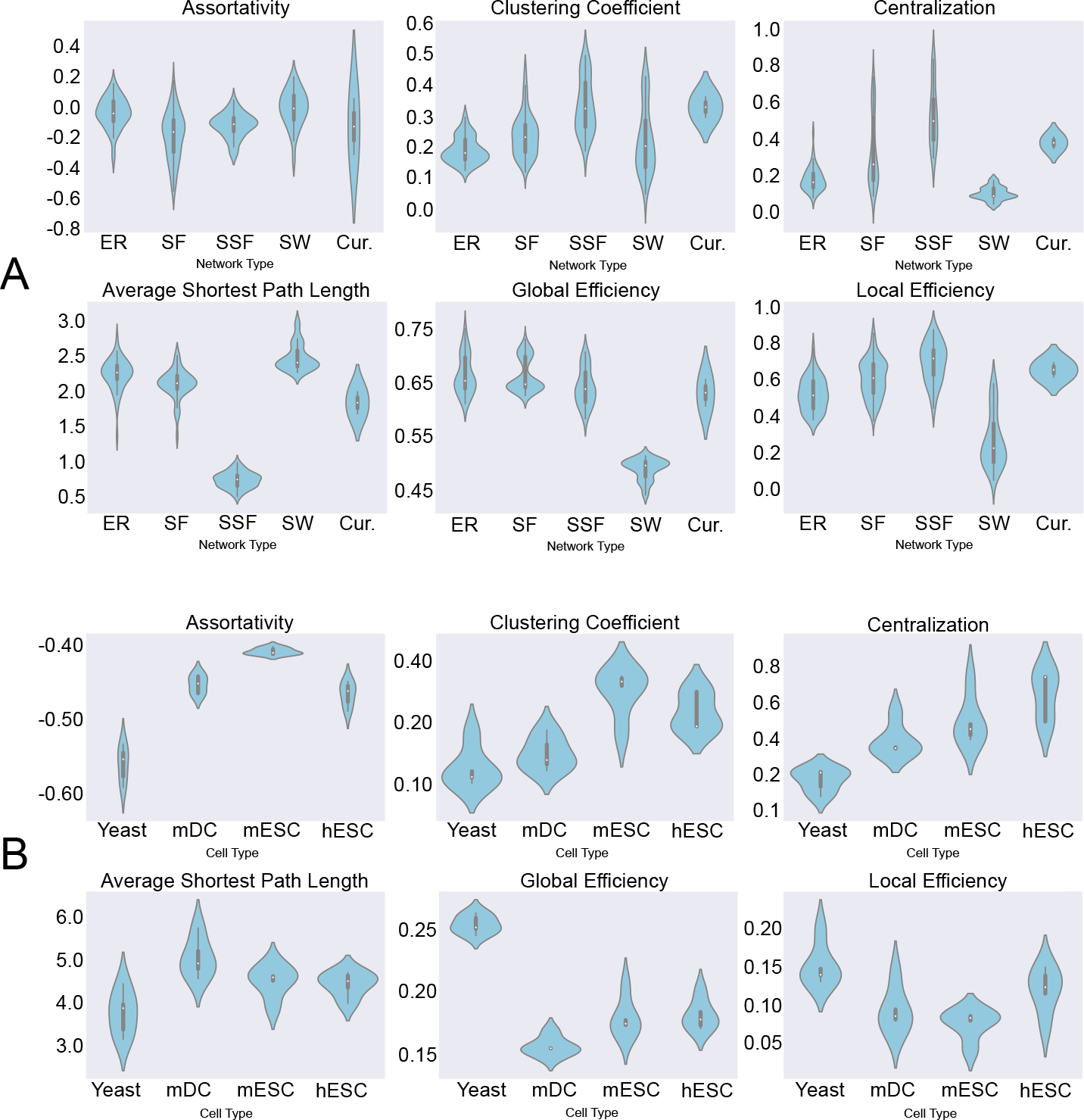
Distribution of the topological properties of the synthetic (panel A) and experimental (panel B) ground truth networks.

## Notes

https://github.com/ScialdoneLab/STREAMLINE

### Competing Interest Statement

The authors have declared no competing interest.

### Summary of Updates

We updated the text of the manuscript

